# Human cortical networks trade communication efficiency for computational reliability

**DOI:** 10.64898/2025.12.11.693716

**Authors:** Kayson Fakhar, Danyal Akarca, Andrea I. Luppi, Stuart Oldham, Fatemeh Hadaeghi, Petra E. Vértes, Ed Bullmore, Claus Hilgetag, Duncan Astle

**Affiliations:** MRC Cognition and Brain Sciences Unit, University of Cambridge, Cambridge, UK; Institute of Computational Neuroscience, University Medical Center Hamburg-Eppendorf, Hamburg, Germany; Department of Electrical and Electronic Engineering, Imperial College London, London, UK; Imperial-X, Imperial College London, London, UK; Centre for Eudaimonia and Human Flourishing, Linacre College and Department of Psychiatry, University of Oxford, Oxford, UK; St. John’s College, University of Cambridge, Cambridge, UK; Montreal Neurological Institute, Montreal, CA; Developmental Imaging, Murdoch Children’s Research Institute, Parkville, Australia; School of Psychological Sciences, The Turner Institute for Brain and Mental Health, Monash University, Clayton, Australia; Department of Psychiatry, University of Cambridge, Cambridge, UK; Department of Health Sciences, Boston University, Boston, MA, USA; Institute of Psychiatry, Psychology & Neuroscience, King’s College London, London, UK

## Abstract

Brains are often described as cost-efficient communication networks, optimally balancing the cost of long connections with the benefits of fast communication. Here, inspired by the “use it or lose it” principle, we present a novel game-theoretic model of self-organizing neural units and show that the brain is, in fact, sub-optimal in both regards: First, we demonstrate that regional competition for connectivity under propagative communication dynamics naturally gives rise to network configurations similar to those derived from the human cortex while being even more efficient and economical. Next, we use a reservoir computing framework to compare the information processing capacity of these networks against those of the brain. Although comparable in performance, the more optimal trade-off comes with a tax on computational reliability. Through synthetic lesions, we show that these networks are fragile because, to optimize for communication, they funnel information through a spatially clustered “oligarchy” comprising a tight set of transmodal hubs. In contrast, the human brain uses a more distributed “rich club” backbone that better resists breakdown following targeted attacks, even when it means higher wiring costs and less efficient communication. This reveals a previously overlooked principle: cortical networks trade both cost and efficiency for *reliable* computation. Thus, our findings highlight computational reliability as another, and even more prominent driver of brain connectivity compared to wiring cost and communication efficiency.

**Teaser:** The brain avoids fragile efficiency: brain networks favor reliable computation, revealing resilience as a hidden design rule.

## Introduction

What principles govern the intricate wiring diagram of the human brain? Aristotle and other natural philosophers, limited in experimental methods, established a top-down view that remained dominant for centuries: *form follows function*. They believed the brain’s ventricles and gyrification pattern to have emerged to support its ultimate function, be it cooling down the body or, as became increasingly apparent later, seating cognitive functions (*1*). However, over a century ago, Ramón y Cajal introduced a radically different idea by proposing a neural-level trade-off instead: *neural units in the brain must minimize costly connections while ensuring efficient communication* (*2*). Networks dominated by short-range connections are cost-effective but suffer from limited communication capacity due to increased intermediate processing steps needed to move information through the network. Conversely, networks with abundant long-range connections benefit from enhanced signaling efficiency but with high metabolic and wiring costs. Thus, the human brain’s complex connectivity pattern, featuring numerous short-range connections and a few selective long-range shortcuts, is thought to represent the optimal balance of this trade-off (*3–5*).

Information flow within large-scale brain networks is constrained because not all regions are directly connected. Cognitive operations rely on efficient exchange of information between distributed neural populations, and cognitive difficulties can arise when that communication is disrupted (*6–11*). Yet it is unclear *what constitutes efficient neural communication.* The conventional approach rooted in graph theory assumes information to flow *exclusively* over the shortest path between two brain regions (*12*), here called *routing* communication dynamics. However, following recent advancements in mapping brain-wide activity patterns, a growing consensus surrounding alternative models of communication is forming. It is becoming clearer that brain regions do not interact this way since routing requires each region to possess complete ‘knowledge’ of the brain’s global network topology (*13*). Efforts to develop more decentralized models of inter-regional communication have broadly focused on *propagation* or *diffusion* dynamics rather than precise routing (*6*, *13*, *14*) (Figure 1). In contrast to routing, propagation assumes information to cascade over all pathways while decaying with distance. Diffusion considers information as a random walker, serendipitously traversing the network until reaching its target. Thus, these alternative frameworks challenge traditional definitions of communication efficiency by assuming communication to unfold over multiple parallel (and longer) pathways instead of just the shortest one.

**Figure 1:**
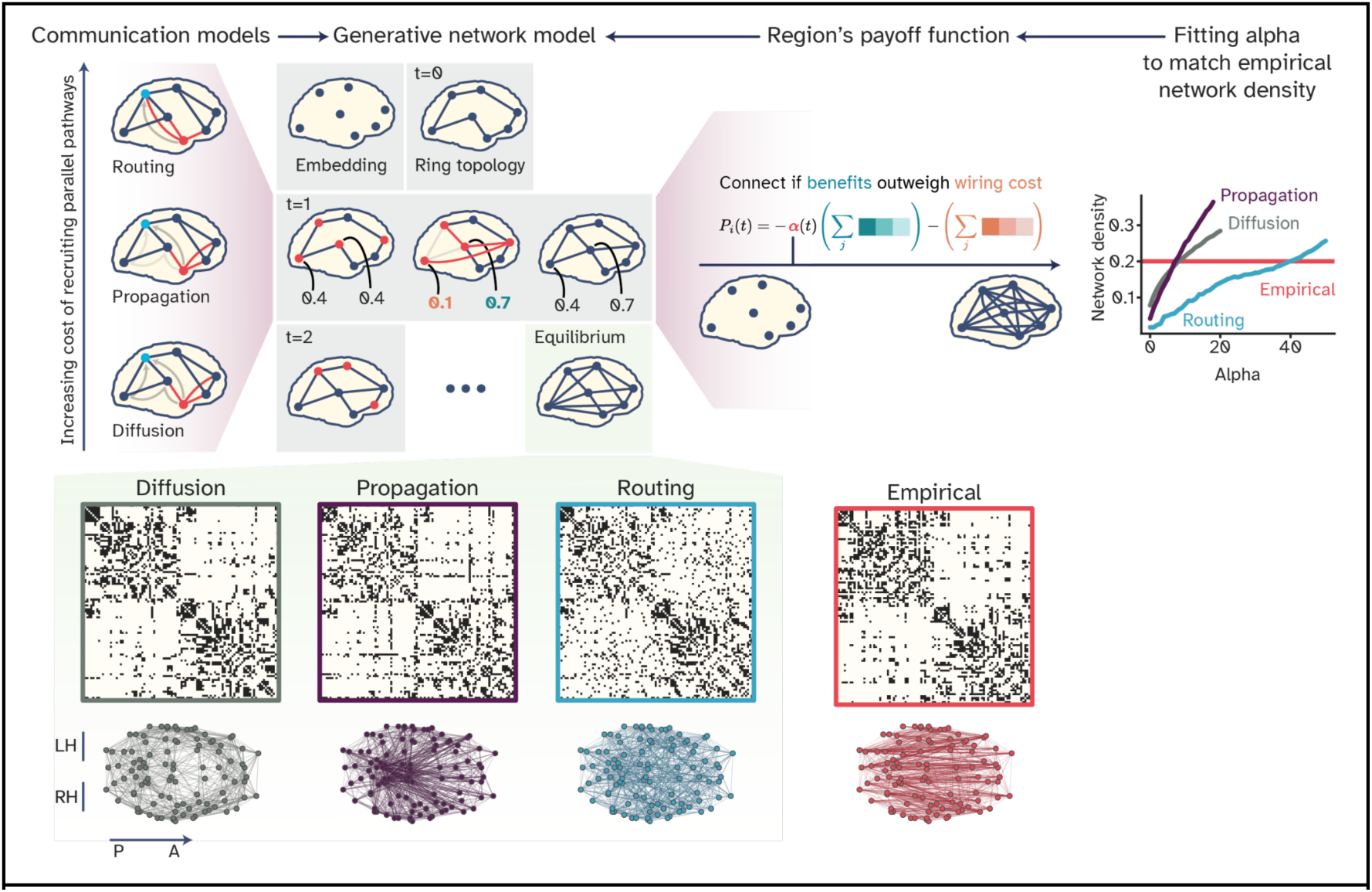
Visual abstract of the framework. At each step of the model, a random group of nodes (red) evaluates its payoff function by comparing the current network configuration with an alternative where existing connections are removed and previously absent ones are added. Each node independently chooses whichever option yields a higher payoff, without considering effects on the rest of the network. The payoff function combines two components: wiring cost, measured as the Euclidean distance between connected nodes, and communication capacity, estimated through one of three models of inter-areal communication. In the **routing** model, capacity depends only on the shortest path length between nodes; in **propagation**, it is based on broadcasting along all available paths, with longer routes contributing less; and in **diffusion**, it reflects the expected number of intermediate steps taken by random walkers. These models represent a spectrum from high to low parallel communication cost, with routing the most expensive, diffusion the least, and propagation in between. A trade-off parameter was then tuned to balance communication efficiency against wiring cost, calibrated to match the connection density observed in empirical brain networks. Note that the opacity of connections in the plotted brain networks is set proportionally to their length to emphasize long-range connectivity, which models disagree the most.

Under these models of neural signaling, could simple and organic processes lead to better network configurations, or ***is the human brain truly at the optimal balancing point of communication efficiency and wiring cost?*** Additionally, ***does better communication naturally lead to better computation,*** or are there other distinct factors determining the brain’s connectivity pattern, e.g., computational capacity? We address these questions by combining game theory, communication models, generative brain network models, and neuromorphic computing. Game theory enables us to formally express the cost-benefit trade-off as a local and distributed optimization problem, where brain regions act as self-interested ‘players’ negotiating connections *without global network knowledge*. We implement these interactions via a generative network model, where regions connect only if the benefit of improved signaling outweighs the wiring cost (Figure 1). Notably, these decisions occur unilaterally, as one region might find a connection advantageous while its target does not, resulting in asymmetric connectivity costs. Finally, we use neuromorphic computing to test the computational properties of networks that result from different trade-offs.

First, brain-like wiring patterns naturally emerged from competitive interactions among self-interested neural elements, each optimizing its own cost-benefit problem. Interestingly, although all emergent networks reflected cortical network characteristics to some extent, only propagation-based networks produced structured long-range connectivity, even surpassing the human brain in terms of both wiring economy and communication efficiency. Incorporating these networks into a reservoir computing framework to solve a memory capacity task demonstrated that human cortical architecture offers superior computational capacity compared to both diffusion- and routing-based but not propagation-based networks. Further analysis of propagation-based networks revealed they optimally navigated the cost-benefit trade-off, primarily by establishing a transmodal ‘high-cost high-capacity backbone for communication’ (*15*) akin to the brain’s rich-club organization. However, this backbone comprised fewer regions and was more spatially localized, resembling an “oligarchy” rather than a distributed rich-club connectivity. Consequently, we show that this centralization introduced both functional and signaling fragility that is averted by the human brain.

Collectively, these findings reveal that local competitive interactions under propagative signaling naturally lead to the formation of complex network architectures, even surpassing the brain in both wiring economy and communication efficiency. However, the brain’s seemingly suboptimal wiring efficiency is offset by its exceptional computational reliability, suggesting an interplay between two evolutionary scales: top-down network-level pressures regulating bottom-up neural competitions.

## Results

To establish a normative benchmark for networks with optimal navigation of the trade-off between wiring cost and communication efficiency, we introduced a novel generative network model grounded in game theory. Briefly, our model begins with a spatially embedded set of regions minimally connected with one another, i.e., a ring graph (but also see Supplementary Figure 2. D). At each time point, a random subset of 10 regions is selected to “play” by disconnecting from their connected neighbors and connecting to the previously unconnected regions within the selected set. Each node then calculates and compares its payoff between the two configurations, opting for the more advantageous one (Figure 1; also see Algorithm 1 in the Methods section). Note that regions do not consider their impact on other nodes, nor do they care about the cost-efficiency of the network as a whole. This process continues until no new configuration is advantageous for any region and a state of equilibrium is reached.

The payoff function was comprised of two elements: the wiring cost of the established links, quantified as the sum of the Euclidean distance between node pairs, and the communication capacity, i.e., total nodal influence derived from three putative communication models. In routing, this capacity was inversely proportional to the shortest path length between nodes, assuming information flows solely through this pathway (*6*). Under diffusion dynamics, signaling is modeled as a random walker, navigating the network serendipitously until it reaches its destination. Here, communication capacity is based on the ‘effective distance’ between pairs, defined as the expected number of intermediate nodes traversed by random walkers between a pair (*16*). Lastly, propagation assumes information to be broadcasted simultaneously along all pathways, with longer ones contributing proportionally less to communication capacity (*17*). Essentially, these models can be arranged along a spectrum of solutions to minimizing the energetic cost of parallel signaling with routing representing the most and diffusion the least expensive parallel communication (Figure 1). A single trade-off parameter α was then adjusted to balance communication efficiency against wiring cost, ensuring the networks have the same number of connections (density) as the empirical data (see Game Theoretical Generative Network Model in Methods).

It is crucial to note that the goal of this framework is *not* to replicate the brain’s connectivity pattern but to create *networks where individual nodes achieve an optimal balance between wiring cost and signaling efficiency*, conditioned on the underlying communication dynamics. Thus, the model remains fundamentally agnostic to the brain’s connectivity pattern, relying solely on the spatial arrangement of nodes and the network’s density. Therefore, the modeling framework may not necessarily converge to brain-like connectivity patterns but to a pattern that optimizes this trade-off. The resulting networks were then used as normative references and compared with empirical data to determine *whether the brain also exhibits characteristics of an optimal communication network*.

Empirical brain networks were constructed from 100 unrelated subjects provided by the Human Connectome Project (HCP) (*18*). These cortical networks were composed of 100 nodes defined based on the Schaefer-100 parcellation scheme (*19*), with a network density of 20%. A group-averaged template of all empirical networks was created to benchmark empirical network similarity measures (Figure 1. A and B, vertical band). Moreover, we constructed four null models: two through rewiring of empirical networks while keeping the degree distribution unchanged (randomized and latticized), one by rewiring the spatially defined long-range connections, and one fully random. These networks represent the extreme configurations of the cost-benefit trade-off. Randomized networks shuffle connectivity while only preserving the nodal distribution of connections, introducing many costly shortcuts. Latticized networks, in contrast, rewire topologically defined long-range connections, forming a web of mainly neighbor-to-neighbor connectivity, all while keeping the degree distribution unchanged. Although this results in networks with long processing pathways, they may still contain spatially defined long-range connections. We addressed this issue by rewiring spatially defined long-range connections (long-range rewired) and by constructing fully random networks with the same density as the empirical network (Erdos-Renyi). To have the same sample size across settings, 100 synthetic networks per condition were generated. All networks were binary and undirected, and reported statistical relationships have p < 0.01 unless specified otherwise.

### Self-organizing networks capture hallmarks of cortical architecture

We used a host of measures to compare generated networks with cortical and null networks, as shown in Figure 2.

**Figure 2:**
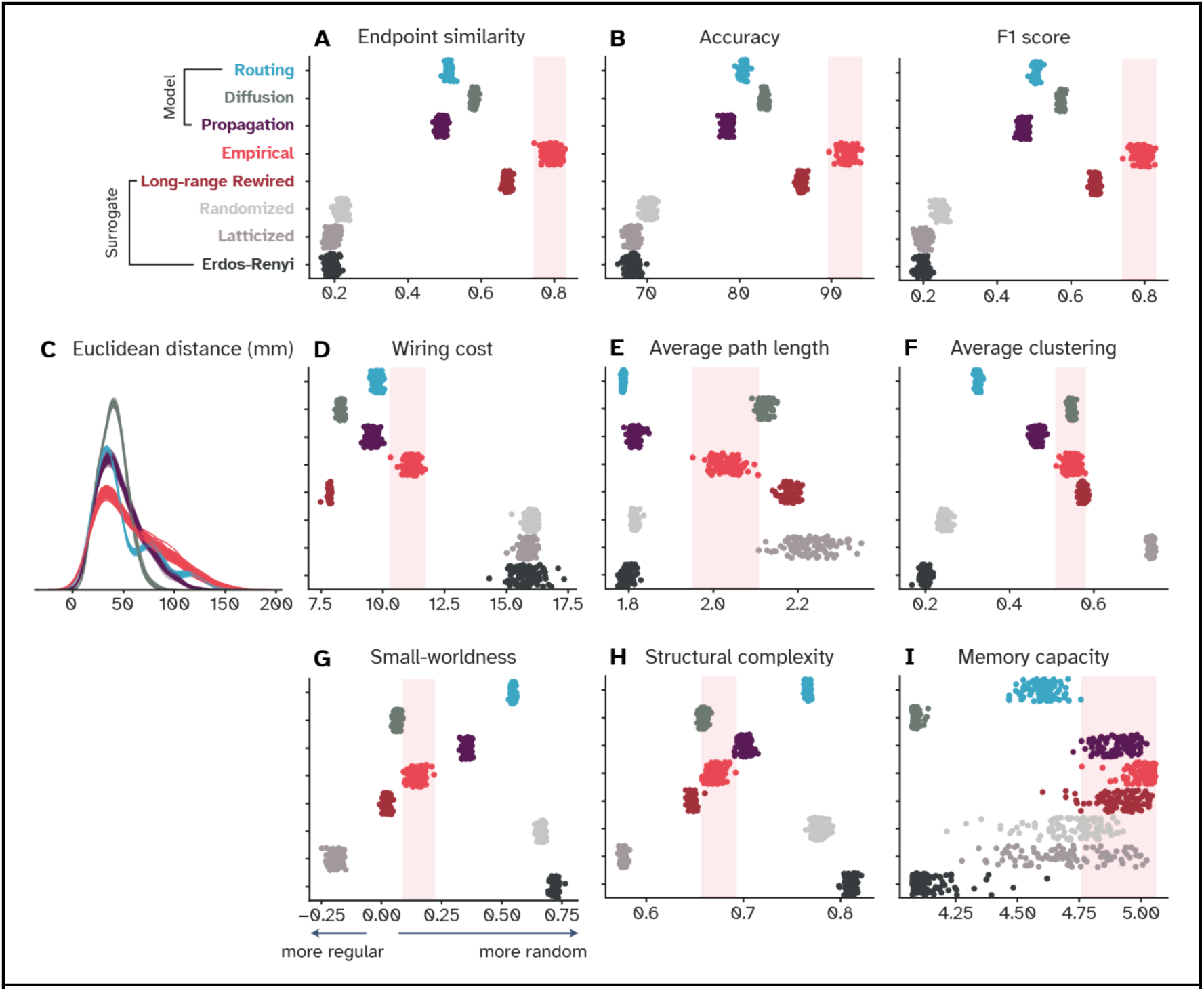
Optimal communication networks capture hallmarks of brain connectivity. Each point represents one network, and the empirical range is shown by the shaded red area. **(A and B)** Model networks capture aspects of empirical cortical wiring evident from the fact that endpoint similarity, accuracy, and F1 scores show better alignment with empirical data than randomized, latticized, or Erdős-Rényi nulls. Individual empirical networks are compared with the group-averaged consensus connectome **(C)** distribution of Euclidean connection lengths across network types. Diffusion-based networks lack long-range connectivity, while routing results in precise wiring that leads to three peaks. Propagation leads to similar long-tail decay but with more short-range and fewer long-range connections compared to empirical data. **(D, E, and F)** Comparison of wiring cost, average path length, and clustering coefficient shows that the human brain is not optimal in either wiring economy or communication efficiency, as traditionally defined based on average path length. **(G and H)** Small-worldness and structural complexity highlight differences in global organization, with diffusion-based models resembling empirical topology despite lacking long-range connectivity. **(I)** Memory capacity results show that empirical networks outperform most generative and null models.

Firstly, we asked ***whether connectivity in the brain shows characteristics of an efficient communication network***. We used ‘endpoint similarity’ defined as the cosine similarity between the connection pattern of the same node in two networks (e.g., emerged from routing-based dynamics vs. the brain) to address this question. A value of one represents the highest similarity, i.e., when a region is linked exactly to all the same regions as in the target network, and no similarity results in zero (see Methods). Averaging across all nodes, we found that networks constrained by all three communication dynamics better capture the empirical data compared to randomized, latticized, and fully random networks (between 0.5 and 0.6 vs. 0.2, Figure 2. A). This is also visible in Supplementary Figure 3A for a comparison based on PCA embedding, which resulted in a similar ranking. Moreover, modeled networks also performed better than null networks in predicting individual links in the empirical brain, given two measures of accuracy (80% vs. 65%) and F1 score (between 0.5-0.6 vs. 0.2, Figure 2. B). Accuracy measures the overall percentage of correct predictions (true positives and negatives), while F1 scores take possible class-imbalance into account by taking the harmonic mean of precision and recall instead. Both metrics show moderate but significantly better alignment, compared to other null networks. Note that the null network generated by rewiring long-range connections is closest in all three metrics, which is expected given that the rest of the network is identical to the empirical data. Also, the highest achievable endpoint similarity and F1 score, measured by comparing the empirical data with a subject-averaged template is about 0.8 and 90% for accuracy. Together, direct comparison of the cortical wiring patterns with those simulated based on connectivity-agnostic self-organizing models showed that the brain contains signatures of an efficient communication network.

To then test ***which topological features of empirical networks are captured by our generated networks***, we measured small-worldness, average clustering coefficient, structural complexity, wiring cost, and average path length. Briefly, average clustering coefficient is a measure of segregation or “cliquishness” in the network. Small-worldness measures the balance of this segregation and global integration in a network. Values around zero show a perfect balance, with negative values representing more clustered and segregated networks and positive values hinting at greater integration resulting from many shortcuts. Average path length measures the expected number of intermediate nodes in processing pathways and is the conventional measure of signaling efficiency under routing dynamics. Structural complexity measures the spread of the network’s eigenspectrum. Larger complexity corresponds to a more diverse distribution of eigenvalues, hinting at a larger pool of possible neural dynamics (*20*, *21*). Together, these plots (Figure 2. E-H) can be interpreted as follows: Networks constrained by diffusion dynamics exhibit brain-like topologies, as they tend to align more closely with empirical values across nearly all measures. For instance, average clustering coefficient of both spans from 0.5 to 0.6. However, these model networks lack long-range connections, suggesting that the high resemblance is driven by the abundance of short-range connections (Figure 2. C and Figure 1). Interestingly, although all three modes of communication result in more cost-efficient wiring patterns compared to the brain, only diffusion results in an average shortest path length that exceeds that of the brain (Figure 2. D, E). This indicates that, according to traditional measures of graph efficiency, diffusion leads to lower communication capacity than the brain due to its lack of long-range shortcuts, even though it precisely mirrors the brain’s average clustering coefficient. Notably, both routing and propagation achieved efficiency levels of random networks while still maintaining more parsimonious wiring cost (Figure 2. E and D).

Lastly, networks derived from routing dynamics tend to exhibit far more randomness than those of the brain (Figure 1 and Figure 2. F and G), with clustering and small-worldness values aligning more closely with random graphs rather than the brain. Collectively, these findings indicate that, firstly, the brain, although still cost-efficient, is not optimally managing either its wiring cost or its communication capacity, as better configurations were achieved. Secondly, constraining diffusion dynamics results in networks lacking long-range connectivity, yet it captures other topological characteristics of the brain better. In contrast, routing produces networks that appear random and fail to replicate the brain’s notable features, such as its average clustering coefficient and small-worldness. Finally, propagation yields more efficient networks, achieving better wiring economy, comparable structural complexity, clustering, and small-worldness to that of the brain.

Each of our simulated networks was optimizing a different communication dynamic, but ***what was the impact of this optimization on that network’s computational properties*?** We evaluated all networks using a neuromorphic computing paradigm known as echo state networks (ESN). These models resemble conventional artificial neural networks, particularly recurrent neural networks, but differ primarily in their training approach (*22*). In ESNs, the hidden layer, or ‘reservoir,’ remains unaltered, while only the readout layer is tuned to solve the task (see Methods, Reservoir computing framework). To estimate the computational capacity of our networks, we used the widely recognized memory capacity (MC) task (*23–26*), where the network receives a sequence of random values as input and must recall lagged versions of it. The further back in the sequence it can reproduce, the better the memory capacity. Previous research with brain-inspired reservoirs has yielded mixed results. While some found brain-inspired networks to outperform random connectivity, others found the opposite (*25–28*). It is therefore unclear whether more efficient communication, as conventionally measured based on routing dynamics and attributed to random connectivity, leads to enhanced computational capacity or not. As we have networks specifically optimized for communication, we could use them to address this open question. If more efficient communication naturally translates to better task performance, then we should expect memory capacity to follow the rough ranking of average path length (Figure 2. E), with networks characterized by shorter average path length performing better. However, memory capacity exhibited a different ranking, with the human brain at the top, followed by propagation-based models and the long-range rewired null model (Figure 2. I). Notably, diffusion-based networks consistently underperformed compared to other networks, including null models. Additionally, fully random networks performed poorly compared to randomized and latticized brain networks (see Figure 2. I, the bottom three categories), suggesting that the brain’s heterogeneous degree distribution, which is also present in these null models, plays a crucial role in computation as well. We replicated these results with a different parcellation based on the Desikan-Killiany atlas with a total of 68 cortical regions (Supplementary Figure 1).

In summary, this section highlights two key findings: first, under propagative signaling dynamics, selfish nodes that optimize their local cost-communication trade-off naturally form networks that exhibit characteristics of human cortical connectivity and computational capacity. Additionally, these models surpass the human brain in terms of wiring economy and communication efficiency, suggesting a suboptimal connectivity pattern in empirical data. Lastly, although the brain outperformed all other networks in our neuromorphic setup, propagation-based networks performed on par with the brain, suggesting a role for structured long-range connectivity in computational capacity. Together, propagation-based networks are more efficient and cost-effective than the brain while also maintaining comparable computational capacity, establishing a relatively *better* wiring configuration. This begs a question: ***could the ‘suboptimal’ cortical connectivity of the human brain be attributed to other evolutionary pressures than wiring economy, communication efficiency, and computational capacity?*** Or is this pattern simply a bad design shaped by phylogenic constraints?

### Human cortical networks prioritize robustness over communication efficiency and wiring economy

Emergence of structured long-range connectivity, cost-effective wiring, comparably high memory capacity, combined with results from previous works (*29–31*) permit us to consolidate propagation as the plausible regime of communication and focus further on the relationship between propagation-based networks and the empirical ones. Doing so, we plotted memory capacity as a function of global propagation efficiency (but see Supplementary Figure 2. A for routing and diffusion) and discovered that the human brain, in fact, resides at the Pareto front of these two objectives (Figure 3. A).

**Figure 3:**
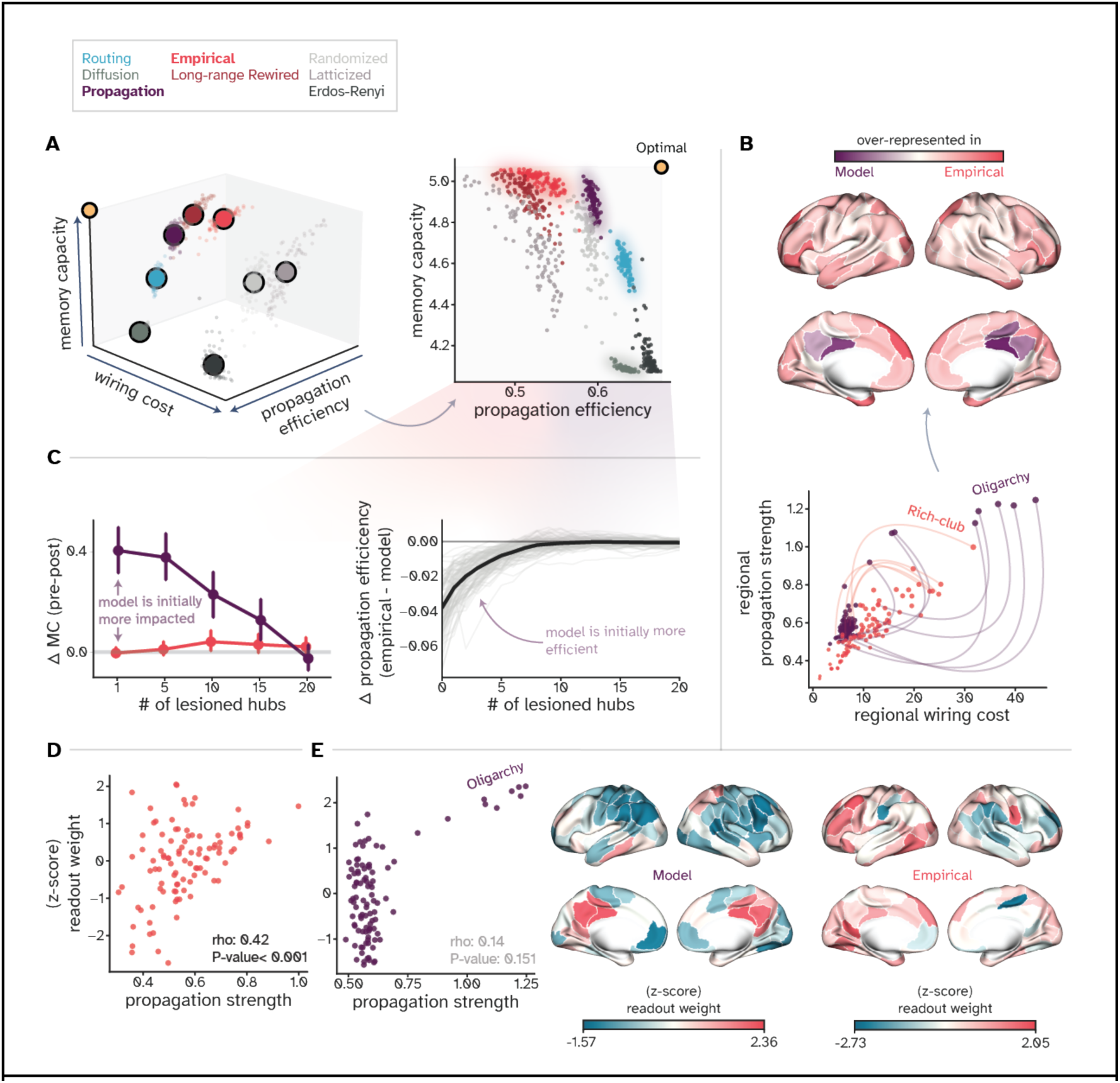
Distributed hub-connectivity makes brain networks less efficient but more robust against targeted attacks. **(A)** Relationships among wiring cost, communication efficiency, and memory capacity reveal a trade-off between communication and computation, with the empirical brain prioritizing robust computation over efficient communication. **(B)** Comparison of regional cost-capacity values shows that the human brain allocates connections in a more distributed fashion, forming a larger set of hubs (rich-club), while propagation-based models converge on a smaller and spatially confined set of oligarchs. Curved lines show discrepancy between high-degree nodes of the two networks, indicating their disagreement in allocating connections. **(C)** Progressive lesioning of hubs in these two networks shows how oligarchy forms a functional bottleneck to enhance information propagation. Empirical networks are less susceptible to performance degradation, while removing even one hub from the oligarchy reduces memory capacity **(D)** In reservoir networks based on empirical connectomes, nodes with stronger propagation strength also show higher functional relevance, measured via their allocated readout weight, while in **(E)** propagation-based networks, this relationship is absent. The functional importance is concentrated within the small oligarchic core, revealing the bottleneck again.

Based on Pareto-optimality theory in evolutionary biology, organisms balance multiple and often conflicting goals, resulting in phenotypes that are not optimal for any single goal (*32*, *33*). Instead, Pareto-optimal solutions are a range of solutions that dominate sub-optimal ones and lie on the ‘Pareto front’ of the solution space, each prioritizing different objectives of the trade-off (*34*, *35*). In our case, an optimal solution unconstrained by a trade-off between memory capacity and propagation efficiency would occupy the upper-right corner of the plot (indicated by the yellow circle). However, all network topologies are confined to a space with Pareto front solutions lying on an arch, suggesting that, after a point, marginal increase in propagation efficiency leads to a rapid decline in memory capacity (see Supplementary Figure 2. A for a clearer visualization). Notably, this arch contains an unstable inflection point between two stable zones of ‘*high performance, low communication efficiency’* and *‘low performance, high communication efficiency.’* On the left-hand side of this zone lies the human brain with reliably high performance while on the right-hand side performance tends to decay as propagation efficiency increases. Said differently, networks resulting from both propagation and routing communication regimes sit on the unstable inflection point, while the human brain networks are spread relatively flat across the high-performance zone, prioritizing reliable computation across multiple instances, i.e., different subjects. This suggests that computational reliability may be a factor driving human cortical architectures away from better communication efficiency, since further increasing communication pushes networks towards the unstable tipping point.

Collectively, (Figure 3. A) suggests that the human brain could further optimize signaling efficiency and still remain at the Pareto front, as propagation-based networks do. However, that neighborhood of the Pareto front appears to be an unstable tipping point, beyond which marginal gains in propagation efficiency leads to a sharp decline in memory capacity. By prioritizing reliable computation, human cortical networks sacrificed both more economical wiring configurations and enhanced communication. This finding aligns with previous research showing that spatially embedded networks perform better and show characteristics of cortical networks when propagation efficiency is minimized as opposed to maximized (*36*, *37*). We further speculate why there might be an inverted U-shaped relationship between communication efficiency and computational capacity in the Discussion section.

Next, we aimed to uncover potential reasons behind the functional instability of the propagation-based networks, i.e., ***why individual differences in instantiation of the same generative mechanisms can lead to wide variation in memory capacity?*** Comparing how these networks allocated resources compared to the brain (Figure 3. B) revealed a critical difference: spatially distributed rich-club organization versus a localized *oligarchy*.

### Rich Club Connectivity as a Design Solution Against Fragile Efficiency

Rich-club organization is a hallmark of brain connectivity across multiple species (*38*, *39*). This organization is formed when well-connected hub regions disproportionately connect to each other, establishing a “high-cost, high-capacity backbone of communication” (*15*). It is proposed that this backbone integrates information across different modalities (*15*, *40*), enhances representational complexity of the network (*41*), and ultimately supports complex cognition (*42*, *43*). Comparing empirical cortical networks with those based on propagation revealed a disagreement in allocating resources, i.e., connections, in the network (Figure 3. B, indicated by the purple and red lines). While both networks possess a spectrum of nodes from low-cost, low-capacity to high-cost and high-capacity regions, they appear to be differently distributed in propagation-based models (Figure 3. B and Supplementary Figure 2. B). Most regions in these models belong to the low-cost, low-capacity category, with a handful of extremely resourceful and influential regions forming a localized oligarchy instead of a spatially distributed rich-club (Supplementary Figure 2. B). These regions are clustered around the posterior cingulate cortex and precuneus, which were previously found to be core members of rich-club organization (*40*). However, in this case, they dominate connectivity, prompting us to ask ***what is the computational role of the oligarchy?*** After all, this pattern of connectivity supports the same level of memory capacity as the empirical brain.

Figure 2. C, D and E answer this question. We used a progressive lesioning scheme of the top 20 regions ranked by degree, tracking memory and communication capacities of the network (Figure 3. C). While the human brain remains functionally robust throughout the lesioning, perturbing the oligarch regions resulted in a drastic reduction of both communication efficiency and memory capacity. Interestingly, the perturbation effect became comparable when the oligarch regions were fully removed, suggesting a re-stabilization of the network after the bottleneck is gone (Figure 2. C). To offer a mechanism for ***why perturbing oligarch regions has such a noticeable effect***, we focused on the assigned readout weights to each region and compared them against their propagation strength (Figure 2. D and E). Since these weights are the only trainable component of our reservoir computing framework, we hypothesized that the network’s hub regions have better representations of the input signal, given their extensive connectivity and capacity to integrate information from a wider range of regions than peripheral nodes. This hypothesis was inspired by the finding that long-range connectivity, specifically among rich-clubs, enhances functional diversity (*41*, *44–46*). We found a robust correlation between nodal propagation strength and readout weight (Figure 2. D) in cortical networks (rho=0.42) but not in propagation-based networks (rho=0.14, p-value=0.15), as the readout layer was predominantly relying on the oligarchy (Figure 2. E). This finding supports our hypothesis and previous research linking hub-connectivity to transformation of information (*41*, *46*). Thus, it becomes clear that oligarchy is the main driver of the relatively high computational capacity seen in propagation-based networks. Consequently, lesioning them degrades task performance since the overall information processing capacity of the network is reduced.

Next, to pinpoint ***how this wiring pattern emerged from local competition for influence***, we performed a cost-benefit analysis for each connection, asking whether the connection justifies its cost for both sides it connects. Briefly, for each edge, we computed the payoff of both nodes, removed the connection, and compared the payoffs with and without the connection. If removing the connection makes one side better off and the other worse, it is likely that the link was established by the region benefiting from it. Interestingly, the results reveal a strong bias against the oligarchy, indicating that this centralized core had emerged due to decisions made by peripheral nodes (Figure 4. A, and Supplementary Figure 2. C). More notably, the spatially proximate neighbors of the oligarchy benefit the most from this arrangement (Figure 4. A). In other words, regions closest to the oligarchy pay very little wiring cost connecting to them while gaining easy access to the whole network via the oligarchy’s extensive connectivity. We then noticed that the topographical organization resulting from this analysis resembles a smooth gradient separating unimodal from transmodal regions (Figure 4. B). Thus, we asked ***whether these peripheral regions are indeed the sensorimotor unimodal areas*** (*47*). Since previous research has shown a disproportionate evolutionary and developmental uptake in wiring of transmodal areas (*48–51*), answering this question could provide further evidence for local and communication-based connectivity optimization to be a key contributor.

**Figure 4:**
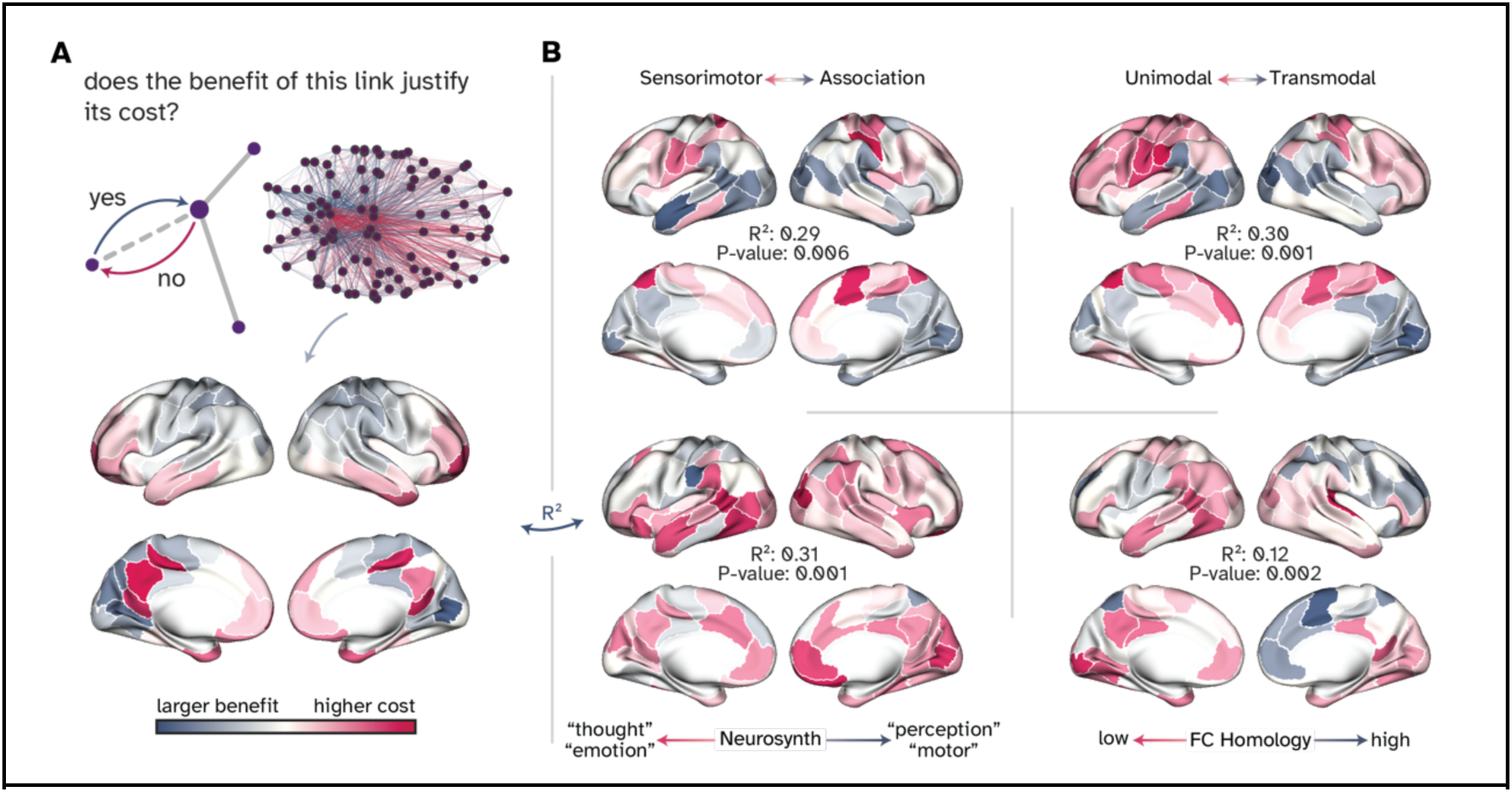
Cost-benefit analysis of individual links and its network-wide organization. **(A)** Cost-benefit analysis of individual connections reveals that oligarchic hubs emerge primarily from peripheral nodes’ aspiration to integrate with the rest of the network without paying the wiring cost. Cortical projection of this cost-benefit analysis highlights that the oligarchy is spatially clustered around posterior midline regions, e.g., Precuneus and posterior cingulate cortex, but also the temporal poles and lateral orbitofrontal cortices. **(B)** These regional differences align with established cortical gradients, including the unimodal-transmodal, sensorimotor-association, large-scale maps of cortical activation during various cognitive tasks (Supplementary Table 1), and cross-species functional homology axes. Reported P-values are adjusted for spatial autocorrelation (see Methods, Null maps for statistical significance)

Therefore, we compared the cortical map of regional cost-benefit values with well-established natural axes of cortical organization: the sensorimotor-association (SA) axis (*48*), the unimodal to transmodal functional hierarchy (*52*), the first principal axis of cortical activation maps during 123 cognitive tasks (Neurosynth) (*53*, *54*), and the cross-species functional homology between humans and macaque monkeys (*55*). Significant correlations in all four comparisons supported the notion that the oligarchy is formed by human-distinct regions of association cortices (Figure 4. B). In essence, the hyper-connected oligarch-club appears to have emerged to enhance communication among, and integrate information from, unimodal regions. However, in the model, and unlike the brain’s rich-club, the circle was further restricted to a smaller set of apex regions to optimize wiring and communication even more. Notably, we also found modest correlations with maps of cortical thickness and gene expression (R^2^=0.16 and 0.27 respectively, P-value=0.02) while finding no relationship with maps of cortical scaling, mean regional blood flow, the second axis of functional gradient, and evolutionary cortical expansion (Supplementary Figure 4). Together, significant correlations with maps related to functional arealization of the brain (sensorimotor-association) alongside nonsignificant correlations with cortical expansion and scaling suggest the following: While connectivity pattern of transmodal regions was, in part, shaped to integrate incoming information from distant unimodal areas, cortical scaling and expansion was driven to meet the required processing demands.

To summarize, results from this section provide an answer for why propagation-based networks are relatively computationally powerful but less robust: optimizing for communication and wiring economy resulted in architectures that naturally develop integrator hub-regions, enhancing their capacity for diversifying representations. However, this organization is also a tight bottleneck, sometimes leading to diminished memory capacity as seen in the previous section (Figure 3. A, C, and D). Moreover, given that oligarch regions are among the brain’s rich-club regions, it could be that, among other factors, external top-down pressure for robustness constrained the enrichment of those regions further into the oligarchy. In other words, the oligarchy is an extreme case of current rich-club organization that can emerge from unrestricted bottom-up local competition for connectivity (Figure 5).

**Figure 5:**
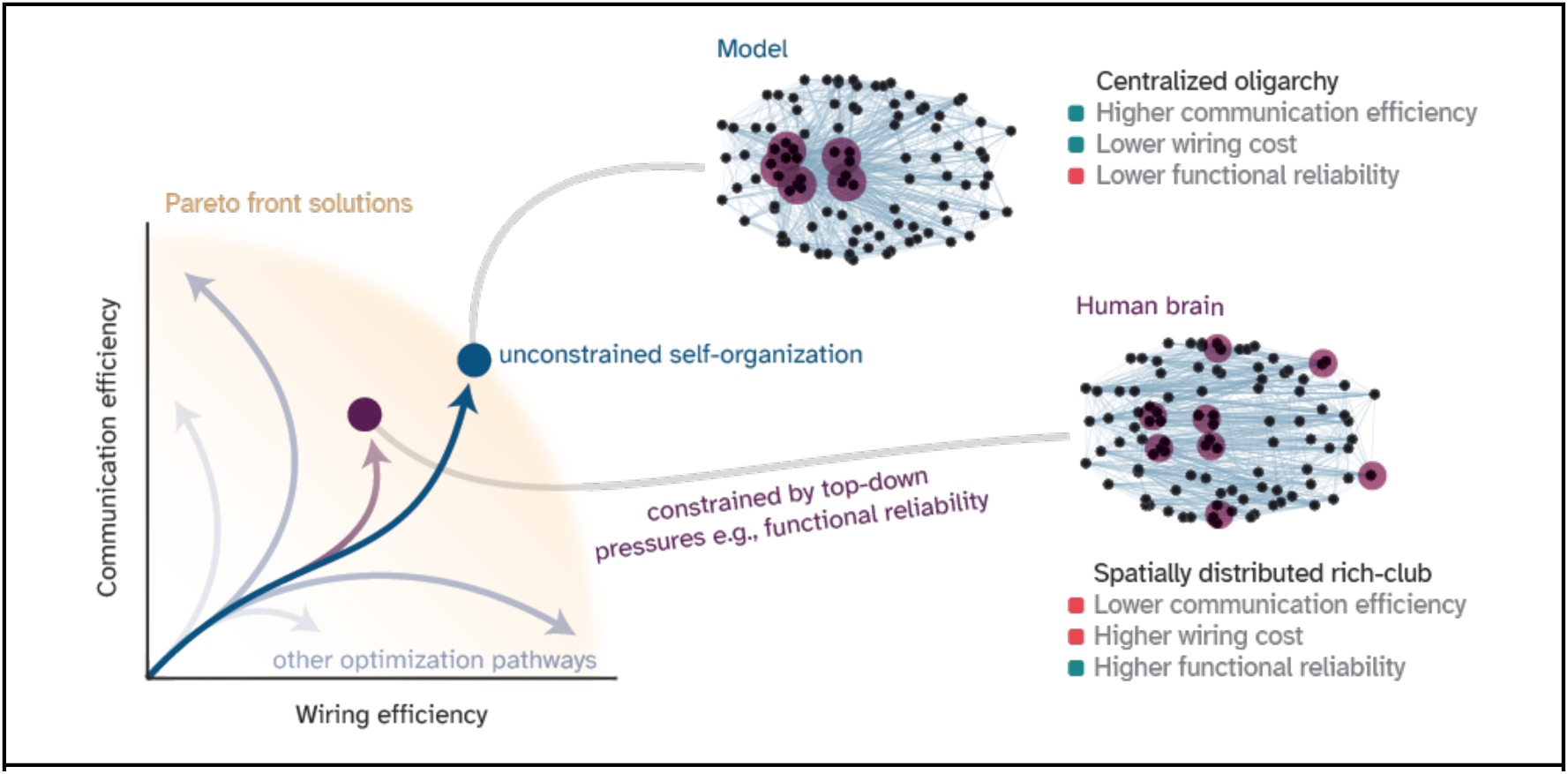
Visual summary of the main findings. While unconstrained local competition for signaling efficiency drives networks toward a hyper-efficient but fragile “oligarchy” (blue), the human brain (purple) settles for sub-optimal configurations that maintain better functional resilience. By adopting a spatially distributed “rich-club” architecture, cortical networks sacrifice better wiring economy and communication efficiency to ensure robust computation, revealing resilience as a critical top-down constraint on brain organization. This schematic is based on the PCA embedding of optimization trajectories shown in Supplementary Figure 3.

## Discussion

We revisited a well-established hypothesis suggesting that a trade-off between communication efficiency and wiring cost is the primary driving force constraining the brain’s complex connectivity pattern. Using a game-theoretical generative network framework, constrained by putative communication models, we discovered that patterns resembling cortical connectivity emerge only under propagative signaling dynamics. Optimal communication networks under routing dynamics place precise long-range connections, resulting in far more random wiring patterns than the brain, while diffusion-based networks contained no long-range connectivity (Figure 2. C). Interestingly, this wiring pattern was previously reported when optimizing for wiring cost by replacing regional positions instead, with the same detrimental effect on communication efficiency (*4*, *5*). Altogether, our findings are consistent with previous research, which indicates that inter-areal communication in brain networks follows a broadcasting-like dynamic, rather than the shortest path routing or random walker-based diffusion (*29*, *30*).

In our modeling framework, individual regions negotiate their own local trade-offs by competing for influence, making the global topology fully emergent. That is, *we only fit the model for the number of connections, not for topological characteristics* as is commonly done (*34*, *56*, *57*). This local competition approach was inspired by the well-studied phenomenon of “use it or lose it”, wherein neural elements compete for resources and can eventually even take over each other’s functions (*58*, *59*). Our results suggest this type of competition also supports the emergence of complex patterns such as those seen in the human cortex. Strikingly, this simple model, constrained by propagation-like signaling dynamics, generated networks that have a better wiring economy than the brain. This design solution was achieved by forming a centralized oligarch-club instead of a distributed rich-club, leading to lower wiring costs, more efficient neural communication, all while maintaining a computational capacity on par with the empirical brain networks. However, we show that the hyperconnected oligarchy is also an information chokepoint. Consequently, these networks concentrate readout weights on just the oligarchy, show more variability in task performance across different instantiations, and degrade faster with synthetic lesions.

Together, these findings add *robust and reliable computational capacity* as another component to the cost-efficiency trade-off, where the human brain invests resources and sacrifices communication efficiency to *consistently* achieve higher computational capacity. Interestingly, a recent work on large-scale recording from the prefrontal cortex and olfactory bulb in developing mice revealed an oligarchic organization where a small set of hub neurons show disproportionate activity (*60*). Strikingly, the study finds that this architecture is preconfigured rather than experience-driven. Together with their findings, our results suggest a key role for neural computation underlying behavior in distributing hubs across the cortex. Further research could model non-behaving model organisms such as brain organoids (*61*) to test whether an oligarchic organization emerges naturally and sustains when there is no external pressure on neural units to coordinate for supporting behavior.

It is important to clarify what our model is not intended to do: based on game theory, this framework is considered normative, i.e., provides a possible answer for *why* the human brain is organized as such and not *how* (*62*, *63*). The difference is that our framework is not equipped to model neurobiological mechanisms by which neural units are assumed to navigate their trade-offs. Our results indicate that maximizing communication could be an evolutionary pressure on neural units, constrained by wiring cost and the dominant regime by which information flows. Based on previous findings (*56*, *64–68*), we hypothesize homophily, i.e., higher preference for connectivity among similar neural units, to be a candidate mechanism implementing the local goal of achieving maximal communication. However, maximizing homophily instead of communication as a local optimization objective result in networks with extreme core-periphery architectures where all nodes link only to their local hubs and not each other (see Supplementary Figure 3. C).

For simplicity and mathematical tractability, our work was limited to binary and symmetric networks. Future work could address both limitations by introducing weights and asymmetric relationships. We expect that introducing an asymmetric wiring cost where no region can impose costs on others to result in a faster convergence (*69*, *70*). Here, we specifically opted out of asymmetric interactions since our networks of interest, i.e., empirical brain data, were also symmetric. Further research using connectomes derived from invasive tract tracing of other animal species could fully embrace the asymmetric version of our framework (*71*). However, introducing weights requires a shift from discrete to continuous games (*72*).

Moreover, we measured memory capacity as a foundational computation that provides a backbone for many other downstream functions. Future work could measure a large array of complementary computations, such as working memory (*73*) or information integration and navigation (*74*) to see whether the expanded trade-off landscape still shows the negative impact of hyper efficient communication. We expect this finding to hold since aside from our result, previous work on information spreading in complex networks also showed an optimal point for signaling efficiency, after which enhanced propagation of information leads to worse outcomes. For instance, works on excitable models revealed that too many shortcuts cause excitation to first quickly spread and then die out (*75–77*), impacting networks’ self-sustainability. It can also lead to rapid synchronization of distant nodes (*41*, *78*), degrading the diversity of information as nodes parrot each other instead of encoding a rich set of features (*79*, *80*).

To summarize our key findings (Figure 5), this work is a proof of principle that, firstly, signaling in brain networks is better modeled using propagative-based dynamics instead of routing or diffusion. Secondly, local and competitive interactions among neural units can lead to the emergence of complex global connectivity patterns, capturing hallmarks of cortical architecture. And lastly, reliable functional capacity is another organizing feature of the brain, leading to a less efficient and costlier rich-club organization as opposed to an oligarchy.

## Material and methods

The dataset used in this work is available at www.humanconnectome.org/study/hcp-young-adult/document/900-subjects-data-release and the following open-source Python libraries were used: YANAT (Yet Another Network Analysis Toolkit) https://kuffmode.github.io/YANAT/ for the game theoretical framework, Echoes https://fabridamicelli.github.io/echoes/ for neuromorphic modeling, Netneurotools https://netneurotools.readthedocs.io/en/latest/ for thresholding empirical networks, Brain Connectivity Toolbox (BCT) https://github.com/aestrivex/bctpy for constructing degree-preserved null models, Networkx https://networkx.org/ for constructing fully random networks, and Neuromaps https://netneurolab.github.io/neuromaps/ for comparing brain maps.

### Neuroimaging dataset and its preprocessing pipeline

We used data from the Human Connectome Project (HCP; (*81*)), restricted to the 100 unrelated subjects from the S900 release. All participants provided informed consent in line with the HCP consortium protocols, and data acquisition was approved by the Washington University’s Institutional Review Board. More detailed information about the imaging protocols and preprocessing pipeline is available at (*82*) and (*83*).

#### Diffusion-weighted MRI acquisition

The diffusion-weighted acquisition protocol is covered in (*84*). Briefly, diffusion MRI scans were collected on a Siemens 3T Skyra scanner using a spin-echo EPI sequence with multiband factor 3. The acquisition parameters were voxel size = 1.25 mm isotropic, TR = 5,500 ms, TE = 89.5 ms. Three diffusion shells were acquired with b-values of 1,000, 2,000, and 3,000 s/mm², each with 90 diffusion directions, plus 6 b0 images.

#### Preprocessing and fiber reconstruction

Minimally preprocessed data from the HCP pipeline (*84*) were used, which included correction for head motion, eddy currents, and susceptibility artifacts. Diffusion-weighted data were reconstructed using q-space diffeomorphic reconstruction (QSDR; (*85*)) in DSI Studio, yielding spin distribution functions in MNI space with 1 mm resolution. Deterministic streamline tractography was then performed with a modified FACT algorithm (*86*) using the following parameters: angular cutoff = 55°, step size = 1.0 mm, minimum streamline length = 10 mm, maximum length = 400 mm, and anisotropy threshold set by Otsu’s method. For each subject, 1,000,000 streamlines were reconstructed.

#### Parcellation atlases and network construction

Structural connectivity was estimated using one primary and another replication cortical atlases. For the primary atlas we used the Schaefer-100 parcellation that defines regional borders based on functional gradients and clustering of resting-state fMRI connectivity (*19*). For the replication atlas (Supplementary figure 2), we employed the anatomically defined Desikan–Killiany atlas, comprising 68 cortical regions (*87*). Streamlines were assigned to regions according to their termination points, and subcortical regions were excluded.

For each subject and parcellation, a structural connectivity matrix was constructed by counting the number of streamlines between each pair of regions. Finally, matrices were binarized after proportional strength-based thresholding to achieve a target density of 20%, consistent with prior studies (*31*, *83*). Specifically, we used an algorithm provided by the python library netneurotools that trims weak connections while preserving a backbone of minimum-spanning tree to ensure the network remains connected. Together, this pipeline produced subject-specific connectomes for 100 subject that formed the empirical basis for comparison with null models and generative networks.

### Game theoretical generative network model

We implemented a game-theoretic generative framework in which each node *i* of the network acts as a self-interested player, locally optimizing a payoff function that maximizes communication capacity given the wiring costs. The network is built first as an embedded ring graph where each node is connected to its two immediate neighbors. The embedding is based on the 3D coordinates corresponding to the centroid of each region. At each iteration, a randomly selected set of 10 node evaluate their payoffs under the current configuration and compares it with the payoff of an alternative configuration where their connections are rewired. Rewiring simply flips the state of the given connection, meaning that, *within the selected set*, it cuts the existing connection and link unconnected nodes. If the alternative yields higher payoff, the node unilaterally updates its connections, i.e., keeps the new configuration. If not, the node defers the new link and keeps the current configuration (Algorithm 1). Since nodes decide unilaterally, new configurations can be imposed on those who did not choose it, as further investigated their implications in our cost-benefit analysis. This procedure is repeated for *T* = 10,000 steps, although an equilibrium state is usually reached within the first 3,000. This equilibrium is visible in (Supplementary Figure 3. A) where the ‘developmental trajectory’ of each model is plotted. It is also noticeable in Supplementary Figure 2. D where we show three random initial networks with different densities eventually settle on the same target density, i.e., the equilibrium. The payoff *U*_*i*_ for node *i* is defined as:

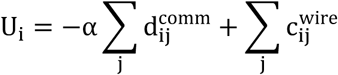

where 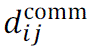 is the communication distance between nodes *i* and *j*, 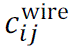 is the Euclidean distance or the wiring cost of the edge. Communication distance is computed across the whole network since the objective is to be as influential as possible. The parameter α is the trade-off parameter controlling the relative weighting of communication over wiring. Effectively, α controls the density of the emerged network and is finetuned using a bisection search with linear interpolation to match the density of the empirical networks as 0.2±0.01 (Supplementary Figure 3. C). The resulting α values are:

|  | Schaefer 100 | Desikan-Killiany |
| --- | --- | --- |
| Routing | 43.01 | 39.25 |
| Diffusion | 8.13 | 4.90 |
| Propagation | 10.46 | 9.79 |

Differences in alpha values can be interpreted as how much long-range connections should be discounted by their added value in efficiency term to be considered. For instance, under a routing dynamics, very long connections are needed to provide enough influence for a node to establish the connections, otherwise not having more connections is preferred that leads to an equilibrium at a sparser network. Lastly, we chose to formalize communication capacity as a cost (see below) so it can be jointly minimized with wiring cost.

#### Algorithm 1

Game-theoretical generative network model

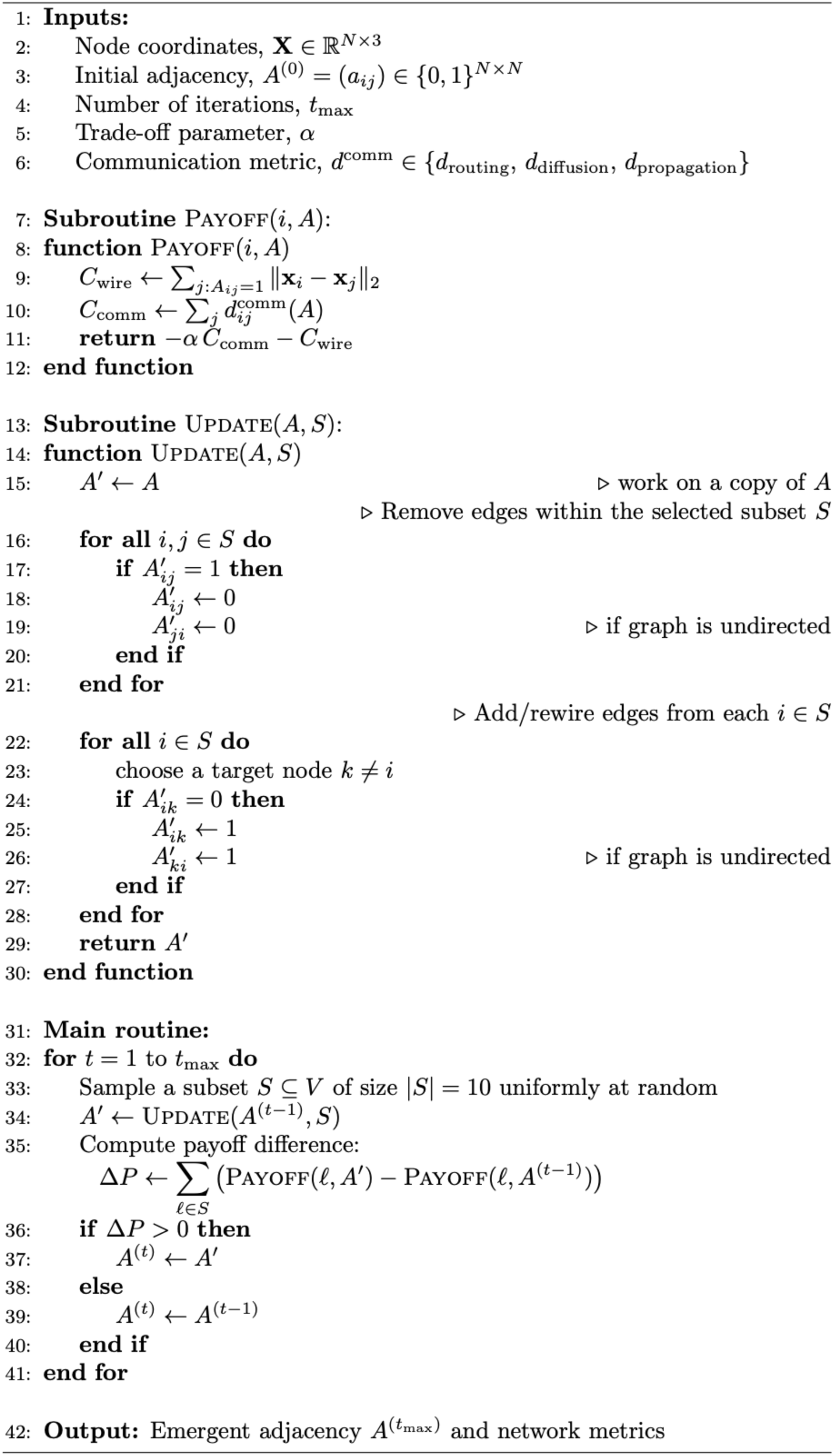

### Communication models

We considered three putative models of interregional communication:

#### Routing (formalized as the shortest-path distance)

Information is assumed to travel exclusively along the shortest path between nodes *i* and *j* using Dijkstra’s algorithm:

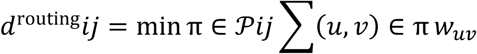

where 𝒫_𝒾𝒿_ is the set of all paths connecting *i* and *j*, and *W*_*uv*_ is the edge weight between nodes *u* and *v*. In this work each edge has unit length, and the shortest path reduces to the minimal number of hops.

#### Diffusion (formalized as the effective distance)

Signaling is modeled as a random walk, where effective distance is proportional to the commute time between two nodes, i.e., the time it takes for a walker to go from one node to the other and returns (*16*). Using the graph Laplacian 𝐿, the resistance distance is defined as:

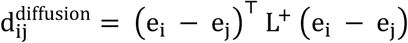

Where, 𝑒*_i_* and 𝑒*_j_* denote the standard basis vectors corresponding to nodes *i* and *j*, respectively, defined as column vectors with a single entry of 1 at the designated index and 0s elsewhere. In this context, the difference (𝑒*_i_* - 𝑒*_j_*) acts as a current operator injecting unit flow at node *i* and extracting it at node *j*. 𝐿^+^ is the Moore–Penrose pseudoinverse of 𝐿.

#### Propagation (formalized as Katz distance)

Information is assumed to broadcast simultaneously over all possible paths, with contributions from longer paths monotonically discounted (*88*). The resulting propagation matrix 𝑃 is computed from

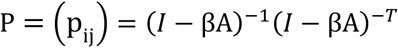

where 𝑊 is the weighted adjacency matrix and 𝛽 is a scaling factor, here set to 0.7, which matches the empirical brain propagation strength (*29*). The communication distance is then

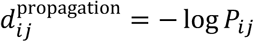

#### Homophily (formalized as topological distance)

As an additional topological constraint, we quantified homophily, defined as the similarity between the connection profiles of nodes (*56*, *65*, *66*) and computed using the matching index of adjacency rows:

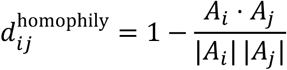

where 𝐴*_i_* is the binary connectivity vector of node *i*. Homophily reflects the tendency of nodes with similar nodal neighborhoods to connect preferentially.

### Null networks

To benchmark our generative networks, we constructed three null models that modify the empirical brain connectome in systematically (*89*). These models are randomized, latticized, and long-range rewired. Additionally, we constructed fully random graphs with the same rough network density (𝑝 = 0.2) as the empirical brain (Supplementary Figure 3. B) using the Erdős–Rényi algorithm.

#### Randomized

Topologically randomized networks were generated by rewiring the original connectivity matrices using the Maslov–Sneppen randomization algorithm (*90*, *91*). This procedure preserves both the network density and the degree distribution, while otherwise randomizing its topology. Concretely, pairs of edges (*i*, *j*) and (*u*, *v*) were selected and swapped such that the new connections became (*i*, *v*) and (*u*, *j*). To ensure sufficient randomization, the rewiring step was repeated 250 times for each connection. Repeating for each connectome, the result is an ensemble of 100 randomized networks that retain certain local connectivity constraints, i.e., the number of connections per region, but lack the global organization of empirical networks.

#### Latticized

Topologically latticized networks were generated using a complementary procedure that biases the Maslov–Sneppen rewiring algorithm towards a ring-like topology while maintaining degree distribution and graph density (*91*). Specifically, swaps of edge pairs were only accepted if the resulting connections placed non-zero entries closer to the main diagonal of the empirical connectivity matrix (which often, but not always, corresponds to connections between spatially adjacent cortical regions). By iteratively favoring such swaps (250 times per connection), the algorithm yields networks in which each node is primarily connected to its nearest topological neighbors, approximating a ring-lattice connectivity pattern. This produces highly ordered connectivity patterns with low global efficiency and high clustering (Figure 2. E and F), providing a structure in contrast with the randomized condition. As with the randomized networks, this process was applied to all 100 empirical connectomes, yielding 100 latticized version.

#### Long-range rewired

In addition to randomized and latticized null models, we implemented a rewiring procedure designed to reduce the wiring cost of empirical networks by selectively swapping long-range connections to isolate their influence on different graph properties depicted in Figure 2.

Starting from the empirical adjacency matrices, we identified all edges whose length (measured via the Euclidean distance) exceeded 70 mm (Figure 2. C). These were classified as “long-range” connections. These edges were then sorted by distance in descending order, such that the longest connections were preferentially targeted.

At each rewiring step (𝑁 = 10), a subset of the remaining long-range edges was removed and replaced with a shorter alternative. Specifically, for a given edge (*i*, *j*) exceeding the distance threshold, the connection was deleted, and each node was evaluated for potential reconnection to its geometrically nearest unconnected neighbor. Whichever node (*i* 𝑜𝑟 *j*) had the closer candidate neighbor was selected, and a new edge was formed. Applied to all empirical connectomes yielded 100 networks with no long-range connectivity. However, other connections were left identical to the corresponding empirical networks.

#### Erdős–Rényi

Lastly, we generated a set of 100 Erdős–Rényi random graphs in which each pair of nodes was randomly connected with independent probability 𝑝 = 0.2, yielding a binomial degree distribution and no higher-order topological constraints. This ensemble of ER graphs serves as a pure random baseline, since while randomization and lattice procedures preserve specific properties of the empirical data (degree distribution or spatial embedding), the ER model establishes expectations for network topology when both constraints, and payoff dynamics are absent.

### Reservoir computing framework

#### Echo state networks

To assess the computational capacity of networks, we implemented them as the core of echo state networks (ESNs) (*22*, *92*). In this setup, the given networks served as the recurrent weight matrices defining reservoir connectivity. The reservoir state was updated according to the equation:

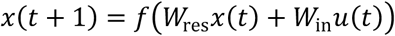

where 𝑥(𝑡) denotes the reservoir state vector at time𝑡, *u*(𝑡) is the external input, 𝑊_res_ is the spectrally normalized reservoir connectivity derived from the underlying network, and 𝑓 is a nonlinear activation function.

For all experiments, we used the hyperbolic tangent (𝑡𝑎𝑛ℎ) as the nonlinear activation function. This choice ensures that the reservoir state trajectories are bounded, while still providing rich nonlinear dynamics. The resulting reservoir states, harvested in response to the temporal input sequences, are later used to train a linear readout that performs the memory task described in the next section.

To enable fair comparison across networks, key parameters were held constant. The spectral radius of 𝑊_res_ was fixed at 0.99, ensuring near-critical dynamics, while input weights 𝑊_in_ were drawn from a uniform distribution in [-1, 1] and scaled by a factor of 10^-5^. Spectral normalization of 𝑊_res_ is essential to ensure the dynamics remain stable and in the echo state regime: without it, the system may become unstable (if the spectral radius exceeds 1) or lose expressivity (if it is too small). This normalization allows the reservoir to retain memory of past inputs while remaining sensitive to new inputs. The scaling of input kept the perturbations small relative to the internal reservoir dynamics, maintaining consistency across different network instantiations. Each network was tested with an identical set of random input sequences (see below). For robustness, we report results as the average performance across 256 independent trials for each network.

#### Memory capacity task and the training paradigm

We quantified reservoir performance using the well-established memory capacity (MC) task (*23–26*, *92*, *93*). At each trial, the network received a one-dimensional signal *u*(𝑡), constructed as an i.i.d. sequence of length 3000 drawn from 𝒩(0,1). The network was trained to reproduce delayed copies of the input, with distinct output units assigned to different delay values τ where τ = {1, 2, …, 40}.

For all lags τ, the readout mechanism, i.e., a linear regression model, was fitted to predict the delayed input *u*(𝑡 − τ) from the current reservoir state 𝑥(𝑡). The fitting solution was done analytically using the pseudoinverse of the reservoir state matrix:

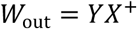

where 𝑋 is the matrix of reservoir states, 𝑌 is the target, where each 𝑌 corresponds to a specific delay τ, and 𝑋^+^ denotes the Moore–Penrose pseudoinverse. Before training, the first 100 transient steps were discarded to wash out initial conditions. No regularization was applied to any of the networks.

A train–test split of 0.8/0.2 was used, with training performed on the first 80% of time points and performance evaluated on the remaining 20%. Memory capacity at each lag was quantified as the squared correlation coefficient ρ between the predicted and target sequences. Finally, the total memory capacity of a network was obtained by summing ρ across all tested lags:

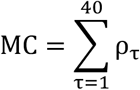

With larger MC values indicating better task performance.

### Network measures

#### Structural complexity

Prior research has shown that large-scale neural signaling is well described as a network of coupled linear systems (*29*, *94*, *95*). This allows us to analyze the magnitudes of eigenvalues λ*_i_* derived from networks’ adjacency matrices to determine the stability and asymptotic regional decay in response to input. A broad distribution of |λ*_i_*| hints at the existence of multiple timescales (*96*, *97*), indicating heterogeneous neural response profiles. Thus, to summarize the network’s capacity for such rich temporal responses, we computed the entropy of its eigenspectrum. Since all our networks were symmetric, the eigenspectrum contains only real parts thus:

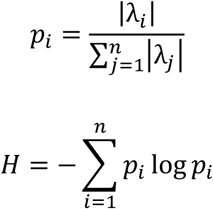

Finally, to normalize values between 0 and 1, we divided by the maximum possible entropy:

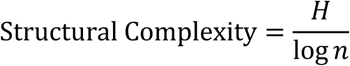

#### Average clustering coefficient

To assess the local segregation of networks, we measured the average clustering coefficient as follows. For each node *i* with degree 𝑘*_i_* ≥ 2, the local clustering coefficient 𝐶*_i_* is defined as the fraction of existing edges among its neighbors relative to the maximum possible number of such edges:

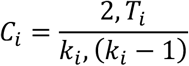

where *T_i_* denotes the number of triangles (three-node closed loops) including node *i*, and 𝑘*_i_*(𝑘*_i_* − 1)/2 is the total number of possible neighbor–neighbor connections. For nodes with 𝑘*_i_* < 2, 𝐶*_i_* is set to zero. The average clustering coefficient of the network is then:

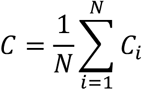

where 𝑁 is the number of nodes. 𝐶 quantifies the tendency of nodes to form tightly interconnected neighborhoods. High 𝐶 indicates the presence of dense local circuits or modules, whereas low 𝐶 implies a more globally integrated but locally sparse topology. In brain networks, elevated clustering is typically interpreted as a signature of localized processing and functional segregation.

#### Average path length

To measure global integration, we computed the average path length of each network using the same shortest-path distances described above in subsection ‘communication models’ under ‘routing’. For any pair of nodes (*i*, *j*), let 𝑑(*i*, *j*) denote the length of the shortest path between them. The network’s average path length is then simply:

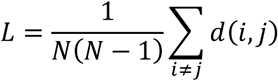

where 𝑁 is the number of nodes. 𝐿 measures the expected number of steps required to connect two arbitrary nodes in the network. Smaller 𝐿 indicates more efficient global integration under the routing signaling dynamics, as information can traverse the network in fewer hops.

#### Small-worldness

To quantify small-world organization, we used the Telesford ω index (*98*), which positions each network on a continuum between lattice and random topologies. For a given network with clustering coefficient 𝐶 and characteristic path length 𝐿, and two sets of degree-constrained ensembles of 100 null networks (described above): (i) a randomized ensemble yielding 𝐿_rand_, and (ii) a latticized ensemble yielding 𝐶_latt_. Together, ω is defined as:

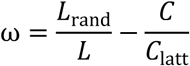

By construction, ω ∈ [−1,1], where values near 0 indicate small-world networks (high clustering like a lattice but short paths like a random graph), ω → −1 indicates lattice-like organization with very high 𝐶 and long 𝐿, and ω → +1 indicates random-like organization with low 𝐶 and short 𝐿. Both these cases are visible in (Figure 2. G).

#### Wiring cost

To quantify the cost of physical embedding for a pattern of connectivity, we computed the wiring cost as the Euclidean length of each edge in three-dimensional space. Node positions were derived from the centroid coordinates of each cortical region in the parcellation. For any edge connecting nodes *i* and *j* with coordinates 𝑟*_i_* = (𝑥*_i_*, 𝑦*_i_*, 𝑧*_i_*) and 𝑟*_j_* = (𝑥*_j_*, 𝑦*_j_*, 𝑧*_j_*), the wiring cost is:

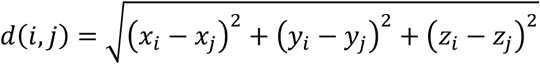

The average wiring cost of the network is the sum of edge lengths across all connections,

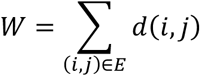

where 𝐸 is the set of edges, and the average wiring cost is obtained by dividing 𝑊 by |𝐸|, the total number of edges.

#### Endpoint similarity

To assess how closely the connectivity patterns of corresponding nodes matched across two networks, we used endpoint similarity. For a given node *i*, we represent its connectivity profile as a binary adjacency vector 𝑎*_i_* (row of the adjacency matrix). Endpoint similarity between networks 𝐴 and 𝐵 for node *i* is then defined as the cosine similarity:

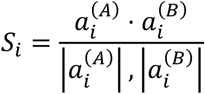

where ⋅ denotes the dot product and |⋅| is the Euclidean norm. Values range from 0 (no overlap in connectivity patterns) to 1 (identical connectivity). The network-level endpoint similarity is obtained by averaging across all nodes:

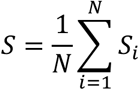

with 𝑁 the number of nodes in the network. Endpoint similarity quantifies the extent to which networks match in their node-wise connectivity profiles, providing a direct comparison of regional wiring patterns between empirical, null, and generative networks.

#### Accuracy and F1 score

To evaluate how well modelled networks predicted the presence or absence of empirical connections, we used two conventional classification metrics: accuracy and F1 score. Given the set of all possible node pairs, predictions fall into four categories: true positives (TP: correctly predicted edges), true negatives (TN: correctly predicted non-edges), false positives (FP: predicted edges that do not exist), and false negatives (FN: missing predictions of existing edges). Accuracy measures the overall proportion of correctly classified pairs:

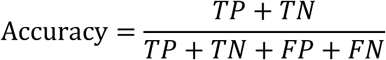

F1 score balances sensitivity and precision by computing the harmonic mean of precision and recall:

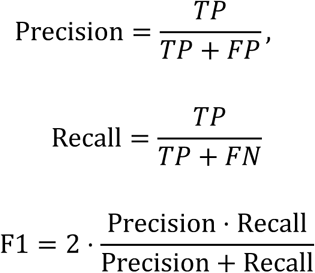

Briefly, accuracy is influenced by the overall class balance (i.e., number of edges vs. non-edges), making it useful for a global measure of prediction quality. In contrast, F1 score is more sensitive to class imbalance, as it emphasizes the trade-off between precision (avoiding false positives) and recall (avoiding false negatives). This makes F1 informative when the number of absent connections greatly exceeds the number of present ones, as is the case in our sparse networks.

### Brain maps

We incorporated cortical brain maps provided through the neuromaps Python library (*99*). Specifically, for the main analysis we used four maps detailed below:

#### Functional connectivity gradient

The unimodal-transmodal gradient was derived from the first functional connectivity gradient introduced by Margulies and colleagues, which methodology is explained in details (*52*). Briefly, the FC1 gradient was derived from resting-state fMRI functional connectivity matrices and obtained using diffusion embedding. This principal gradient is anchored at one end by primary unimodal regions (visual, somatosensory/motor, auditory) and at the other end by transmodal association regions, most prominently the default-mode network (DMN). Intermediate values correspond to heteromodal areas that integrate across sensory modalities (*52*).

#### Sensorimotor-Association gradient

The second gradient was the sensorimotor–association (SA) gradient from Sydnor et al. (*48*) and provides a unidimensional ordering of cortical regions from primary sensorimotor cortex to transmodal association cortex. This axis is constructed by rank-ordering parcels along multiple cortical features and averaging those ranks, yielding an archetypal gradient in which lower values index sensorimotor-like cortex and higher values index association-like cortex. Detailed methods and validation are provided in the source paper (*48*), however, briefly, Sydnor et al. derived the archetypal SA axis by aggregating parcel rankings from ten heterogeneous data types (e.g., T1w/T2w myelin, FC principal gradient, evolutionary expansion, cortical thickness, aerobic glycolysis, receptor/transcriptomic proxies, etc.) computed on the 180 left-hemisphere parcels of the HCP multimodal atlas, then averaging these ranks to produce a single cortex-wide gradient that was used here.

#### Functional connectivity homology gradient

The third gradient was the functional connectivity homology between human and macaque monkey (*55*). This map quantifies cross-species similarity of cortical functional organization between humans and macaques. Briefly, FC homology is built from a joint functional space via diffusion embedding of a concatenated similarity matrix. FC homology at each human vertex is computed as the maximum cosine similarity of the first set of gradients (top ∼15) within a 12 mm searchlight around the surface matched location in macaque, yielding a vertexwise index of functional homology. As reported in the source (*55*), FC homology is highest in unimodal systems (e.g., early visual cortex including V1/MT, auditory, somatomotor) and lowest in transmodal association cortex, with minima in default-mode network territories (e.g., angular gyrus, posterior medial cortex, lateral temporal cortex), consistent with an evolutionary hierarchy from sensorimotor to association regions.

#### Neurosynth activation map of meta-analytic cognitive tasks

Lastly, to contextualize our findings with cognitive functions, we used the cortical map corresponding to the dominant principal component of Neurosynth terms that overlap with the Cognitive Atlas. This map was generated by Yarkoni et al.’s Neurosynth framework (*53*), which automatically aggregates neuroimaging studies to produce whole-brain association maps for psychological terms. Poldrack et al.’s Cognitive Atlas provides a curated ontology of cognitive constructs (*54*). The intersection of these resources yields 123 cognitive terms (e.g., attention, pain, working memory, social cognition; full list in Supplementary Table 1), each associated with a meta-analytic cortical activation pattern. The first principal component of these maps captures the dominant axis of variation, and the resulting cortical map assigns each region a score capturing its loading on this axis, situating them along an axis dominated by perceptual/motor tasks vs emotion and social (*48*, *100*).

#### Null maps for statistical significance

Lastly, to assess the statistical significance of correlations between cortical surface maps while controlling for spatial autocorrelation and hemispheric symmetry, we used the spherical rotation (spin) test, also available in Neuromaps (*99*, *101*). In brief, each cortical map was projected to a sphere, and then random 3D rotations were applied to the spherical coordinates to generate null maps that preserve spatial contiguity and smoothness but disrupt anatomical alignment. Rotations are applied identically within a hemisphere and with sign-constrained axes across hemispheres to preserve contralateral symmetry; vertices rotated off the cortex (e.g., into midline/corpus callosum gaps) are omitted or reassigned by nearest-neighbor on the rotated sphere. The empirical correspondence statistic is compared to its permutation null formed by repeated spins. For each comparison, we performed 𝑁 = 1000 independent spins. At each iteration 𝑘, a random rotation matrix 𝑅*_k_* (uniform on 𝑆𝑂(3) with hemisphere constraints) was applied to 𝑋 to obtain 𝑋^(*k*)^, and the Pearson’s correlation coefficient was recomputed. The (two-sided) spin-based p-value was then estimated as

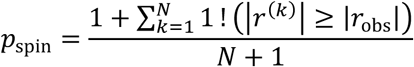

yielding inference under the spatial-permutation framework and controlling inflated false positives that arise from naive parametric tests on autocorrelated surfaces.

## Acknowledgment

Funding for this work was generously provided by the Templeton World Charity Foundation, Inc under grant TWCF-2022–30510 (KF, DAK, DAS, PV, EB) and Deutsche Forschungsgemeinschaft (DFG) SPP 2041/GO 2888/2-2 (KF, CCH).

All research at the Department of Psychiatry in the University of Cambridge is supported by the NIHR Cambridge Biomedical Research Centre (NIHR203312) and the NIHR Applied Research Collaboration East of England. The views expressed are those of the author(s) and not necessarily those of the NIHR or the Department of Health and Social Care

## Author contributions

Conceptualization: KF, DAS

Methodology: KF, DAK, FH, SO, AIL

Investigation: KF

Resources: AIL

Visualization: KF

Supervision: DAS

Funding acquisition: CH, DAS, DAK, PV, EB

Writing—original draft: KF, DAS, FH

Writing—review & editing: KF, DAK, AIL, SO, FH, PV, EB, CH, DAS

## Supplementary figures

**Supplementary Figure 1:**
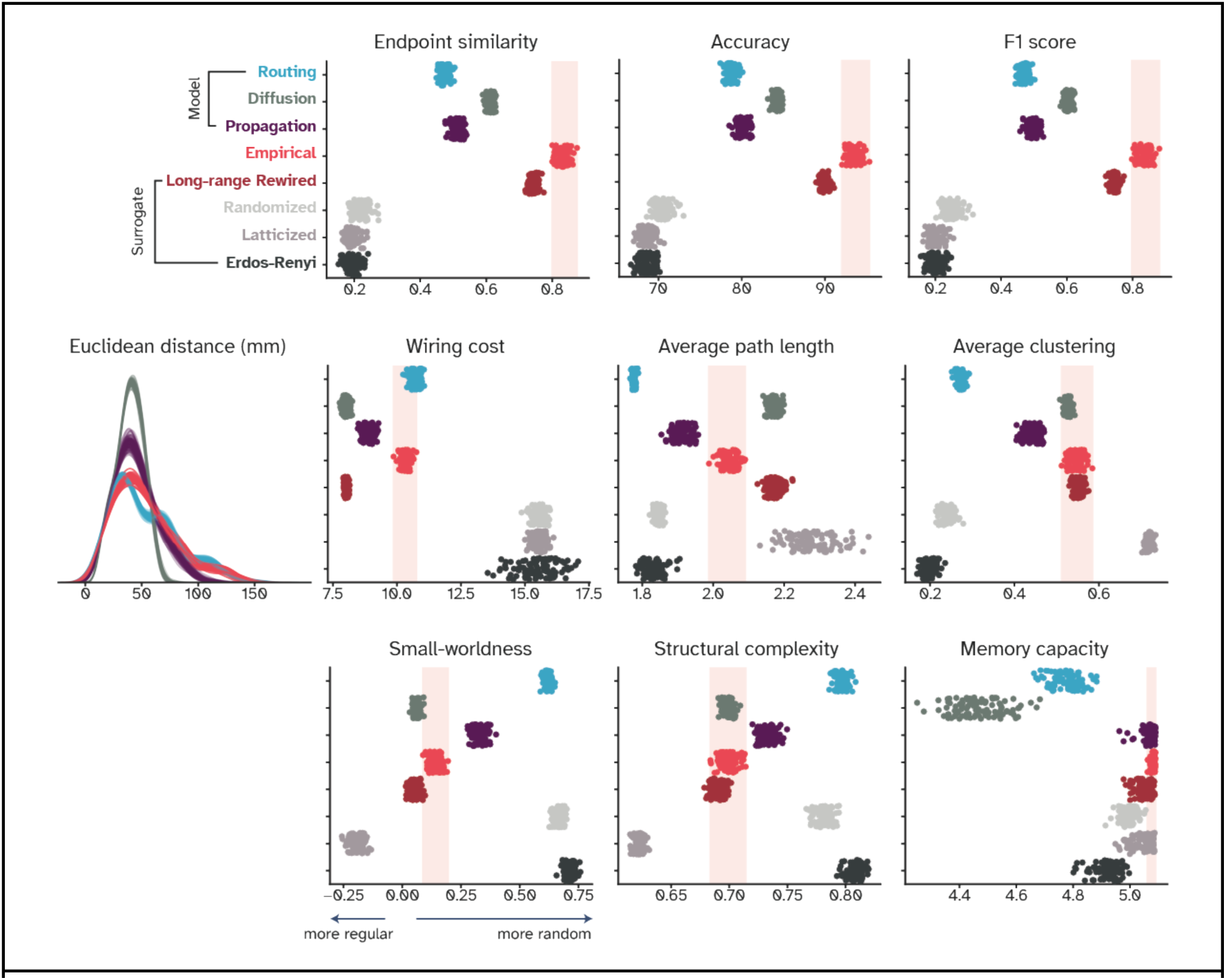
Replicating the main findings using Desikan-Killiany atlas of cortical parcellation.

**Supplementary Figure 2:**
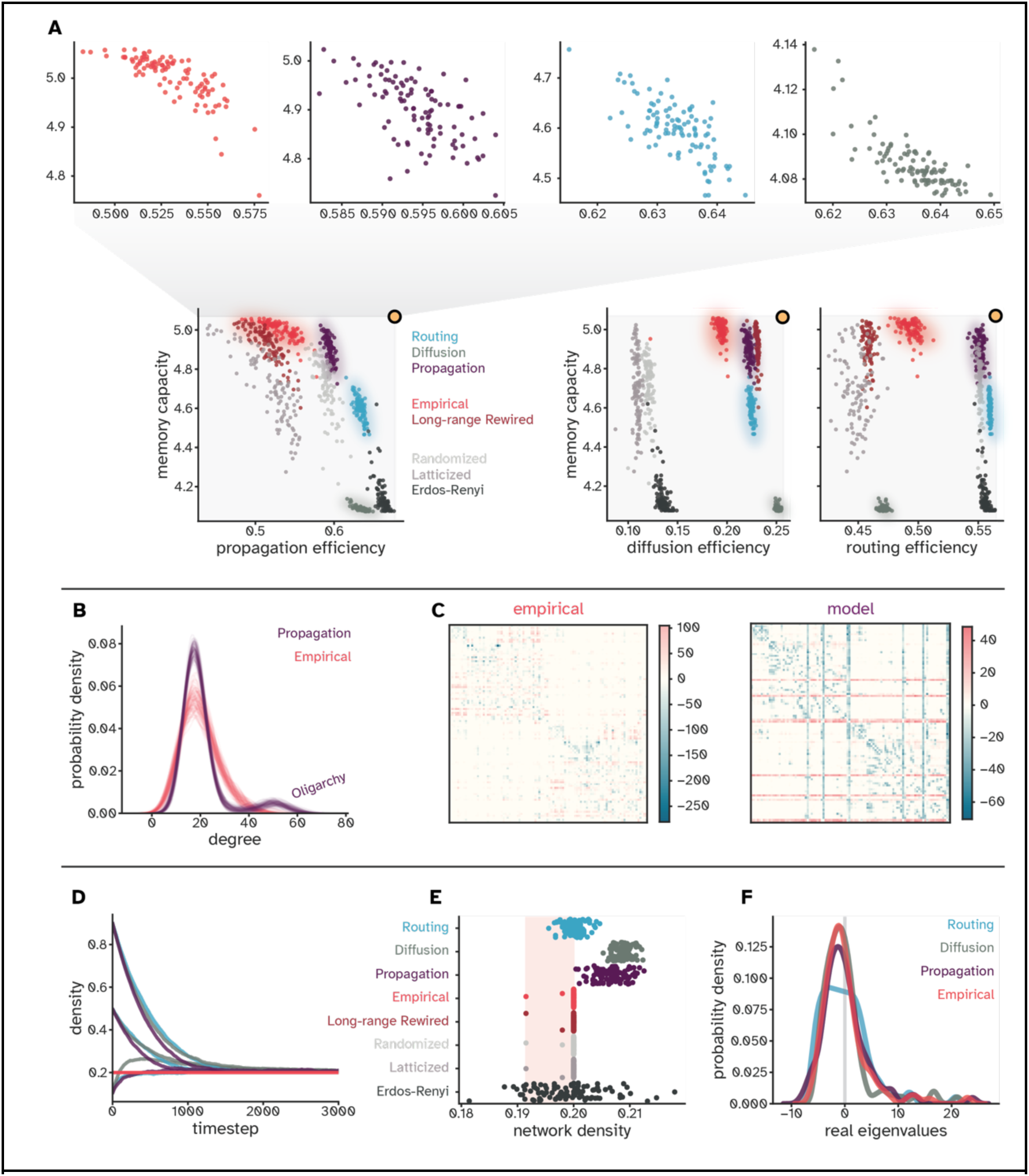
Extended analyses of model networks. (A) Relationship between memory capacity and efficiency measures under different communication regimes. The human brain (red) lies on the Pareto front regardless. (B) Degree distributions show the emergence of a hyper-connected oligarchy in propagation-based models compared to the empirical cortex. (C) Cost-benefit maps show how model networks asymmetrically concentrate resources into oligarchic hubs, unlike the more distributed empirical pattern. (D) Impact of lesioning hub regions on task performance of the ESN. Lesioning was performed by ranking regions by degree and setting input weights of target regions to zero. Values are the difference between task performance prior and after perturbation, thus, positive values mean performance was better pre-lesion. (E) Impact of lesioning hub regions on communication efficiency of networks. The values represent the difference between empirical efficiencies and modelled ones. Thus, positive values represent better post-lesion communication in the brain compared to the model and vice versa. (F) Developmental trajectories of network density across different random initialization reveal convergence toward a stable equilibrium. (G) Final network densities closely match empirical targets with ±0.01 error margin. (H) Spread of real eigenvalues that is measured in structural complexity for empirical and model networks.

**Supplementary Figure 3:**
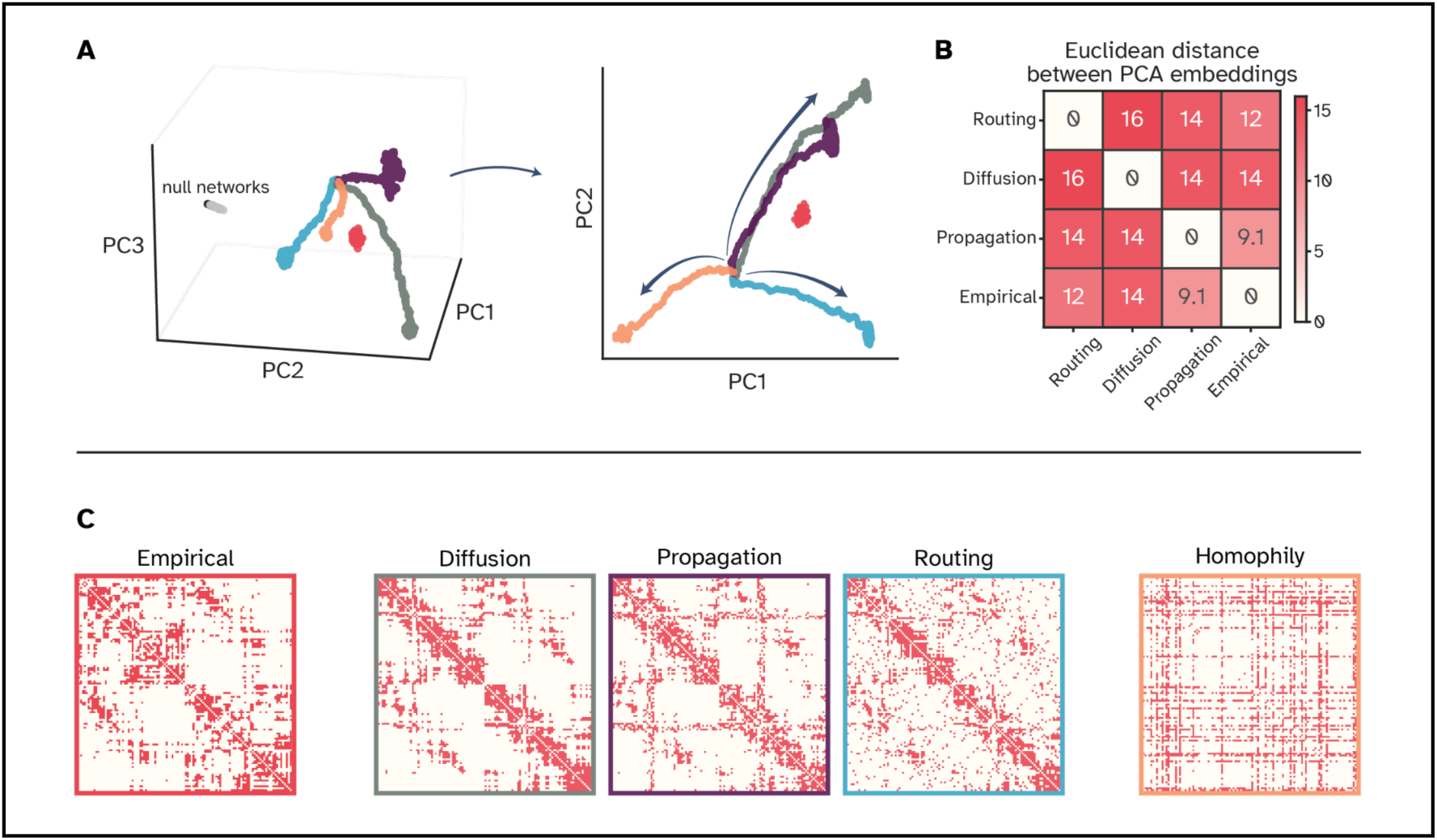
PCA-based embeddings of networks. **(A)** Joint PCA embedding of networks shows the developmental trajectories they take and their settle points. **(B)** Pairwise Euclidean distances between these points quantify the relative separation of network types, showing closer alignment of propagation-based networks with empirical data. **(C)** Connectivity matrices illustrate wiring patterns of empirical, diffusion-, propagation-, and routing-based networks, compared with a homophily-driven model. Homophily produces an extreme core–periphery structure, distinct from both empirical and communication-based networks.

**Supplementary Figure 4:**
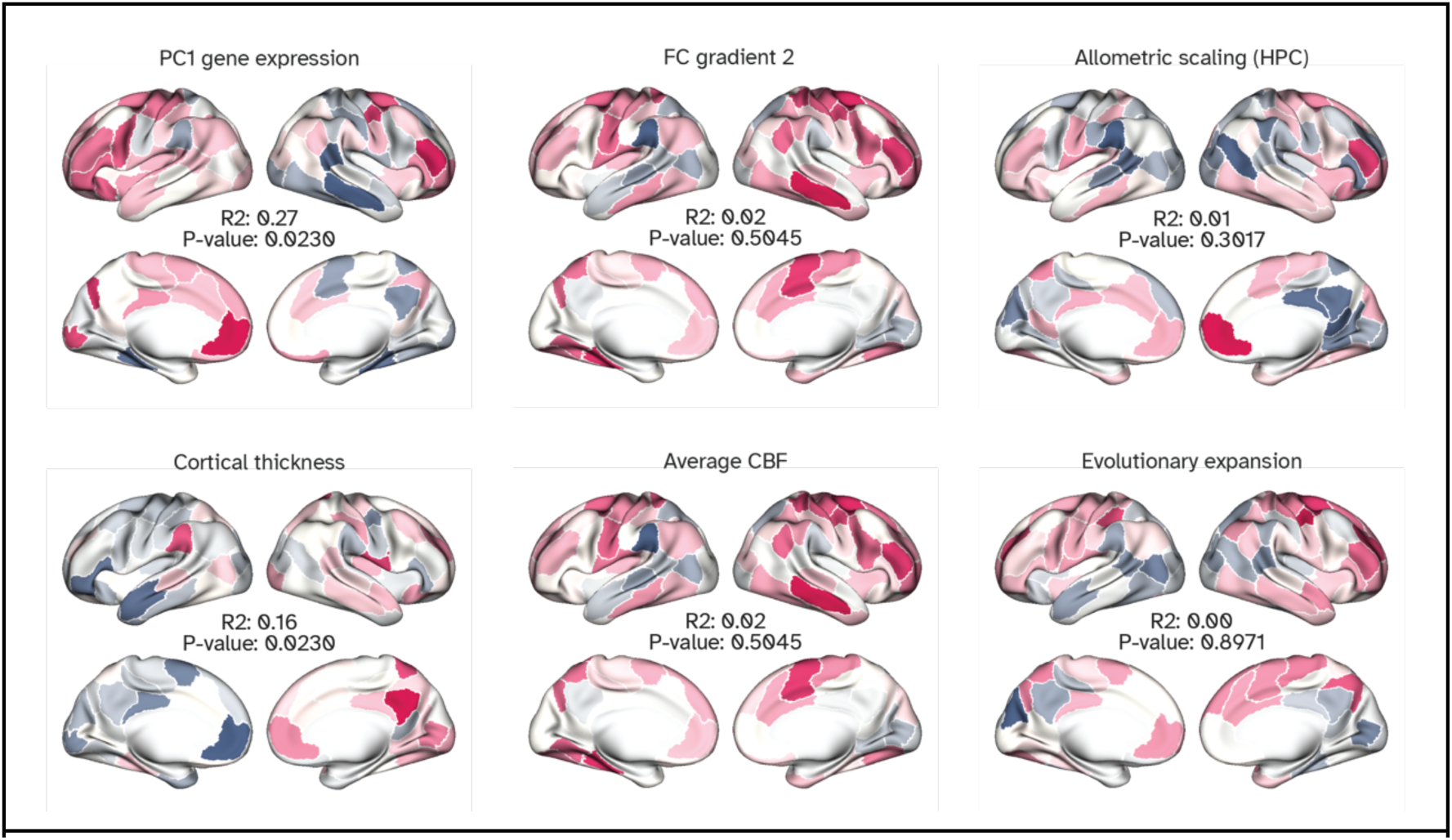
Comparison of payoff dynamics with other cortical maps. These maps are explained in detail elsewhere ***(55)***. However, briefly, the gene expression map is constructed from the first principal component of the Allen Institute’s Human Brain atlas ***(102)***. The second gradient of FC is from **(52)**, the allometric scaling represents the scaling of individual regions in a healthy population (***103***), map of cortical thickness is from ***(104)***, average CBF represents the regional cerebral blood perfusion and is from (***102***). Lastly, map of evolutionary expansion is from ***(55)***.

**Supplementary Table 1:**
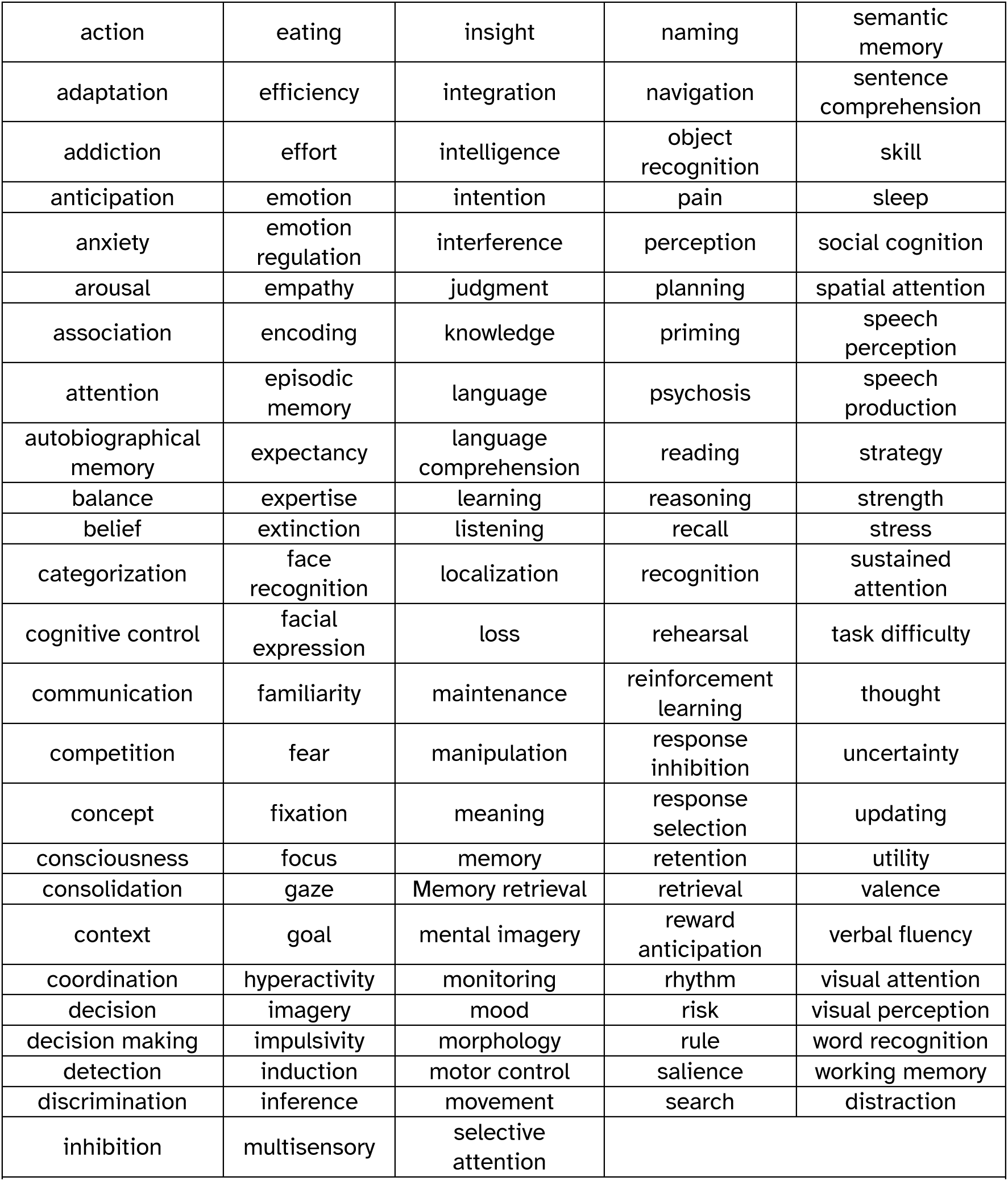
List of 123 cognitive tasks that are used in the Neurosynth PC1 map *(100)*.

|  |  |  |  |  |
| --- | --- | --- | --- | --- |
| action | eating | insight | naming | semantic memory |
| adaptation | efficiency | integration | navigation | sentence comprehension |
| addiction | effort | intelligence | object recognition | skill |
| anticipation | emotion | intention | pain | sleep |
| anxiety | emotion regulation | interference | perception | social cognition |
| arousal | empathy | judgment | planning | spatial attention |
| association | encoding | knowledge | priming | speech perception |
| attention | episodic memory | language | psychosis | speech production |
| autobiographical memory | expectancy | language comprehension | reading | strategy |
| balance | expertise | learning | reasoning | strength |
| belief | extinction | listening | recall | stress |
| categorization | face recognition | localization | recognition | sustained attention |
| cognitive control | facial expression | loss | rehearsal | task difficulty |
| communication | familiarity | maintenance | reinforcement learning | thought |
| competition | fear | manipulation | response inhibition | uncertainty |
| concept | fixation | meaning | response selection | updating |
| consciousness | focus | memory | retention | utility |
| consolidation | gaze | Memory retrieval | retrieval | valence |
| context | goal | mental imagery | reward anticipation | verbal fluency |
| coordination | hyperactivity | monitoring | rhythm | visual attention |
| decision | imagery | mood | risk | visual perception |
| decision making | impulsivity | morphology | rule | word recognition |
| detection | induction | motor control | salience | working memory |
| discrimination | inference | movement | search | distraction |
| inhibition | multisensory | selective attention |  |  |

